## Supplemental materials for "Shaping the developing homunculus: the roles of deprivation and compensatory behaviour in sensory remapping"

### Supplementary Results

#### Activity analysis

To complement the peak position analysis, we also analysed activity levels at the peak of each individual participant's body part representations along S1. Specifically, when comparing individual Z values for peak activity we identify significant main effects of limb difference for the closer neighbours of the deprived hand – the residual arm and the lower face showing stronger activity in CLD individuals (arm: $F_{1, 226.50} = 71.70$, $p < 0.001$; lower face: $F_{1, 226.77} = 14.32$, $p < 0.001$), but not in further body parts (feet, legs and torso; all $p > 0.179$). This indicates that, while deprivation impact the distribution of the body parts along the extent of the homunculus, activity levels are affected more locally. Interestingly, the closest neighbour to the deprived hand area—the upper face—did not show a main effect of limb difference ($F_{1, 276.31} = 2.33$, $p = 0.128$). This result, which replicated a similar effect (Root et al., 2022), indicates that the increased activity found for the deprived hand's other neighbours may not be independent of use.

Age effects were mostly anecdotal (a main effect for the lower face: $F_{1, 226.77} = 7.62$, $p = 0.006$ and the feet: $F_{1, 213.11} = 12.03$, $p = 0.001$); a weak trend towards a main effect of age for arm ($F_{1, 226.50} = 2.84$, $p = 0.093$, was further confirmed by a Bayes analysis: $BF_{10} = 8.18e^{6}$). Interactions between age and limb difference were also sparce, and only conclusively apparent with Bayes statistics (arm: $F_{1, 226.50} = 3.11$, $p = 0.079$, $BF_{10} = 5.84e^{6}$; feet: $F_{1,213.11} = 3.72$, $p = 0.055$, $BF_{10} = 83.52$). This spatially intermitted pattern of results diverges from the more localised effects found for deprivation, and indicates that S1 activity and remapping may further scale with age, providing further evidence for the opportunity for the features of the body maps to be modulated later in life. Nevertheless, as activity levels are more dependant on the stimulation paradigm and strength (in comparison to peak location), we did not follow up on this initial result. For the complete results of the omnibus ANOVA (across the 5 main body parts) see Table S2-8.

#### Computational Model

In this section, we conducted a parameter sensitivity analysis, a method used to assess how variations in a model's input parameters influence its output (Magosso et al., 2010; Manabe & Wetherald, 1967; Straka et al., 2022), to explore the role of behaviour in homunculus adaptation and the model's robustness to parameter changes.

##### *Role of behaviour for homunculus adaptation*

In the Results section of the main article, we used the same *stimulation probabilities* for both the CTR and CLD participants, meaning that behavioural differences between the groups were not necessary to explain the observations using the homeostatic model. Here, we examine the potential influence of behavioural differences between the two groups.

The exact stimulation probabilities for individual body parts, which would encompass the full range of human activities (e.g., sleeping, walking, etc.), are not well established, even for the CTR group. Since the main differences in body part usage between CLD and CTR participants likely occur during manipulation tasks, we used our behavioural data to identify which body parts are used significantly more or less by the CLD group compared to CTR participants. Our data show that the CLD group uses the leg, torso, and lower face more frequently, while the arm is used less (see Figure 1B). However, because our behavioural data mainly reflect manipulation tasks, these values are not directly applicable to the homeostatic model, which also needs to account for non-manipulative activities. Therefore, we tested the model under different sets of body part usage probabilities to capture a broader range of activities.

We hypothesised that increased stimulation of certain body parts in CLD individuals could counteract or even reverse the effects of hand stimulus deprivation, particularly in areas closely linked to the more frequently stimulated parts. For example, before homeostatic adaptation, if hand deprivation reduces a neuron's mean activity by -0.5, this reduction triggers an increase in synaptic weights once homeostasis is applied. Now, in a different scenario where another body part is more often stimulated, resulting in a +0.7 increase in mean neural activity, the combined effect on the neuron would be +0.2 (0.7 - 0.5). In this case, homeostatic adaptation would lower synaptic weights to restore the neuron's activity to its baseline level. This adjustment would be in the opposite direction of the changes seen with hand deprivation alone. In summary, we predict that increased use of certain body parts in CLD individuals could mitigate or even reverse cortical changes linked to hand stimulus deprivation.

To qualitatively analyse the impact of more frequent leg use in CLD individuals, we performed a parameter sensitivity analysis by adjusting the stimulation probability for the leg in the CLD group while keeping the CTR group's probabilities unchanged from the baseline. Specifically, we increased the leg's stimulation probability by 25%, 100%, and 200%, with all other model parameters remaining as in the baseline scenario. **Figure S10** compares the original model with these modified versions.

The model produced two distinct outcomes based on the level of leg stimulation increase. With a 25% increase (i.e., 125% of the baseline), none of the key observations were violated (see **Figure S10**, Panel B). However, with a 100% or 200% increase, observations 2 and 3 were violated. Notably, at a 200% increase, the magnitude of the peaks for the foot and leg were higher for CTR participants, deviating from the expected pattern (see **Figure S10**, Panel D). Additionally, the expected peak position for foot activity was also disrupted.

Similarly, we examined how the homunculus adapts when there is a decrease in arm and hand nerves stimulation in CLD individuals. We reduced the original stimulation probability by 15% (which corresponds to the decrease observed in our behavioural data during manipulation tasks) and by 30%. Since it is unlikely that overall daily activity would decrease by more than double the observed reduction during manipulation tasks, we did not explore reductions beyond 30%. **Figure S11** presents a comparison of all conditions for arm and hand nerve stimulation.

In the lower stimulation scenario, the peak magnitudes for CLD participants were higher, reinforcing Observations 1 and 3, with none of the key observations being violated. Additionally, for CLD participants, there were slight shifts in the activation peaks for torso and upper face toward the centre of the homunculus strip, consistent with Observation 2. This indicates that reduced limb-related stimulation due to behavioural changes in CLD participants has effects similar to reductions caused by limb difference alone.

In summary, guided by behavioural data, we systematically investigated the effects of increased leg stimulation and decreased upper limb stimulation in CLD individuals. For the leg, a 25% increase did not violate any of the four key findings but increases above 100% violated Observations 2 and 3. The results suggest that the empirically observed higher body part usage in CLDs does not generally contribute to the crucial observed activity differences between both groups. Instead, it appears to reduce these differences. However, these differences remain present unless the behavioural differences exceed a certain threshold. For reduced upper limb stimulation, decreases of up to 30% did not violate any of the four key observations. The decrease even strengthens some of the activity differences between both groups suggesting that it has a similar effect on the activity patterns as the input reduction caused by limb difference alone.

##### *Rudimentary scenario homeostatic model*

To demonstrate the robustness of our model results to variations in parameter choices, we present a simplified, or "rudimentary," experimental scenario. Unlike the base model, this version assumes no input from the palm and finger nerves in CLD individuals, meaning there is no peripheral reorganization. Additionally, for both CLD and CTR groups, all body parts—including the palm and fingers—are assigned identical stimulation probabilities ($P_{BP}$=0.05 for all body parts), and the same bell-shaped initial connection patterns between body parts and the homunculus strip are used (see Figure S12).

Figure S13 shows node activity during tactile stimulation of individual body parts. Remarkably, the model replicates most of the key empirical findings (see Results section): 1) The arm activation peak in the CLD group is higher and shifted to the right compared to the CTR group; 2) except for the foot, all CLD activation peaks are shifted toward the centre of the strip relative to CTR peaks; 3) all activation peaks in the CLD group are higher than those in the CTR group. However, in this simplified scenario, empirical property 4—where there is a decrease in activation peak magnitude from the foot to the leg and torso, and an increase from the upper face to the lower face—is only reproduced in the CTR group, not in the CLD group.

In summary, even in this rudimentary scenario, the model replicates most of the key empirical findings, highlighting the robustness of our results. However, to fully capture all the empirical observations, a more complex model is required—one that incorporates peripheral reorganization, higher hand (or corresponding nerve) stimulation frequencies, and stronger initial connections between the CTR hand and the cortical strip.

#### Topographical changes beyond S1

We also examined activity changes in secondary somatosensory cortex (S2), to determine if the changes found in S1 are cascaded downstream. We repeated the same set of analyses reported in the S1 hand area (see Results section in the main text). When looking at activity levels across body parts and groups (Figure S14), we observe a significant interaction effect between age and body part ($F_{4,248}$ =7.08, p<0.001), which was driven by the arm and chin (arm: $F_{1,135.17}$=4.28, p= 0.040; chin: $F_{1,135.17}$=8.82, p=0.004). We found a trending significant main effect of limb difference for the chin only ($F_{1,135.17}$=3.68, p= 0.057), which remained ambiguous following Bayes analysis ($BF_{10}$=0.97), and no significant interactions with limb difference, both for the omnibus model or the follow up ANOVAs per body part (all p > 0.101). When considering representational dissimilarity, we again find no effects of limb difference (all p>0.177) and we observed a significant main effect of age ($F_{1,62}$=5.18, p= 0.026) as adults showed larger distances than children.

To further confirm and assess the representational similarity in S2, we also performed a correlation analysis between the pairwise distance values in the S1 deprived hand area and in S2 for each participant. All groups showed a significant correlation between the two ROIs as indicated by one-sample t-tests against zero (all p<0.046), confirming that the representational structure is overall shared across ROIs. We observed no main effect of limb difference ($F_{1,63}$ = 0.07, p = 0.792), but we observed a significant main effect of age ($F_{1,63}$ = 6.28, p = 0.015) and a weak (and ambiguous) trend towards an interaction ($F_{1,63}$ = 2.951, p = 0.093, $BF_{10}$=0.92), which was driven by larger correlations in control adults relative to control children ($t_{63}$=3.03, p=0.004). Together, these findings indicate that the differential activity observed along S1 to express a limb difference may not be read out by S2, though null results should be interpreted with caution.

### Supplementary Figures

#### Figure S1: Proportion of time analysis across sub-tasks.

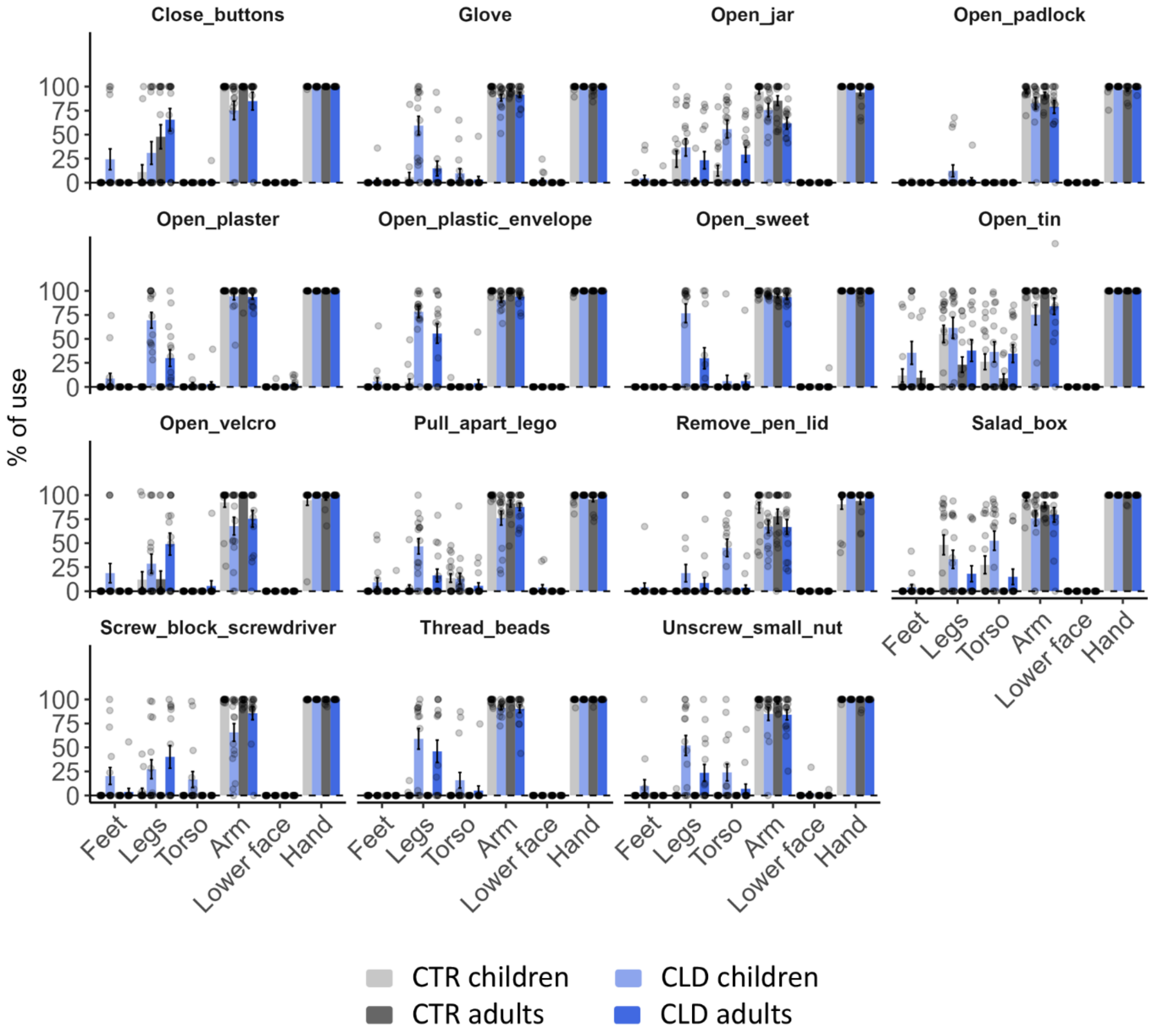


***Figure S1.*** **Proportion of time each body part was used relative to the total task time for each task.** Performance data for each of the 15 tasks used to index compensatory behaviour (for the complete list of the tasks use, see Table S1). Individuals in the CLD group (represented by blue bars), and children in particular (light blue) tend to rely on alternative body parts to compensate for impaired function across a range of tasks that simulate everyday activities. For example, when performing tasks such as putting on a glove, individuals with CLD, especially children, often use their legs. In tasks requiring greater power grip, such as opening a salad box or a tin, the compensatory actions become even more variable. Here, individuals in the CLD group frequently utilise their feet, legs, and torso, with children exhibiting the most diverse range of adaptations. These findings underscore the flexibility and adaptability of compensatory strategies in CLD individuals, with children showing a heightened reliance on alternative body parts during task performance.

#### Figure S2: Absolute time analysis for behavioural task.

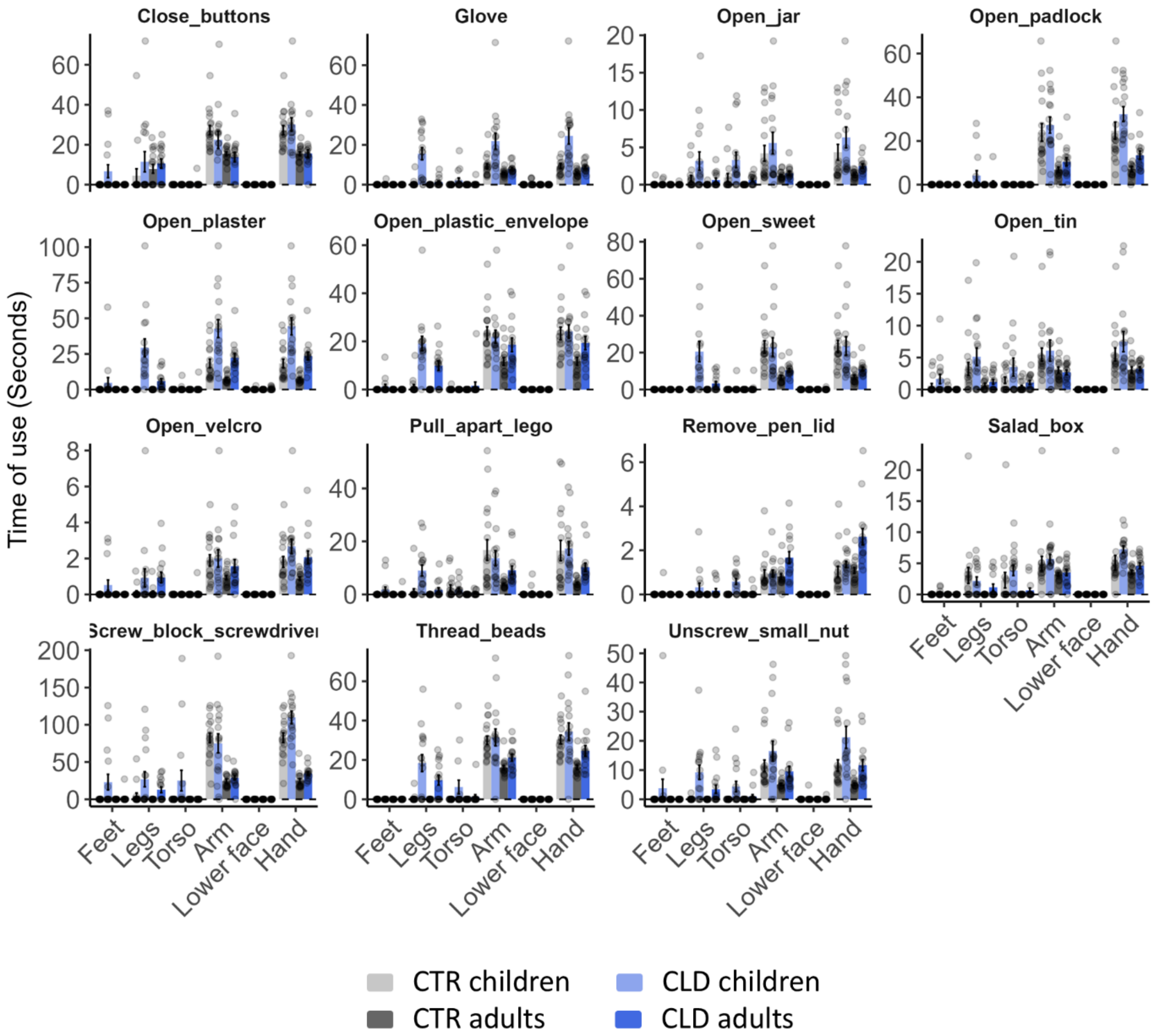


*Figure S2.* **Absolute time in seconds, during which each body part was used in each task.** The time taken to complete the various tasks differed, reflecting the varying levels of task complexity. Adults (represented by darker-coloured bars), with no particular differences between CLD group (dark blue) and the CRT group (dark grey), generally completed the tasks more quickly than children, suggesting that the dexterity required for these tasks is not fully developed in younger participants. This difference in performance is most pronounced in the 'Screwdriver' task (see Table S1), which emerged as the most challenging task for children.

#### Figure S3: Global S1 ROI

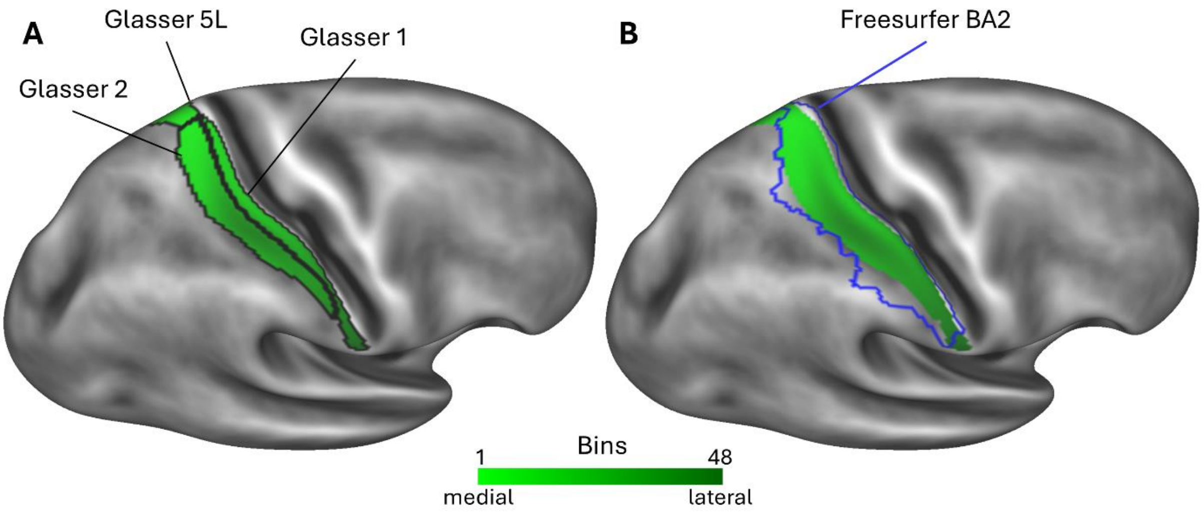


***Figure S3. S1 ROI along the post-central gyrus.*** *To investigate global changes across all body parts along the entire postcentral gyrus, we defined an ROI in each hemisphere, visualised using gradients of green in the plots. This ROI, which included the Glasser's labels 1, 2, and 5L (Glasser et al., 2016), was mapped onto the standard flat maps (FS_LR 32k) of the Human Connectome Project (HCP). The right hemisphere is displayed as an example. The shaded gradients of green represent the number of bins and their orientation along the medial-to-lateral axis. Panel A illustrates the bins-ROI along with the outlines of the labels from Glasser's atlas (Glasser et al., 2016), which were used to delineate the ROI. For comparison, Panel B provides the un-thresholded outline of the FreeSurfer BA2 label.*

#### Figure S4: Activation maps

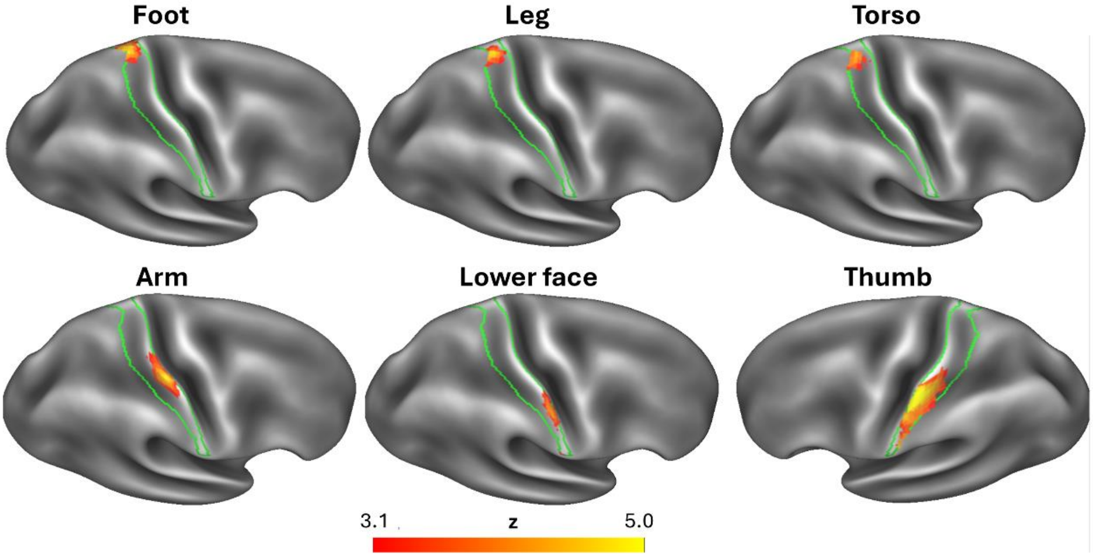


***Figure S4. Activity maps when the different body parts are stimulated****. Group analysis, comprising all study participants, shows the activity patterns (z values) for each body part, reflecting the typical topographic organization (see the classical S1 topography in Figure 4 of the main manuscript) and along the higher-order sections of the somatosensory BA1-2 strip (highlighted in light green). As expected, lower body parts (Foot, Leg, Torso) elicited activity in the medial section of S1, while arm-related activity was observed in a more central region of S1. The upper body, specifically the lower face, activated the most lateral part of S1. All these activities were recorded in the hemisphere ipsilateral to the missing or non-dominant hand, corresponding to the deprived/dominant hemisphere. Stimulation of the thumb of the intact/dominant hand (in CLD/CTR participants, respectively) generated localised activity in the contralateral, intact/dominant hemisphere, as expected, in a well-established location (please note that the thumb establishes the lateral boundary of the hand representation). These findings support the consistent somatotopic organization of S1. FMRI data processing was carried out using FEAT (FMRI Expert Analysis Tool) Version 6.00, part of FSL (FMRIB's Software Library, www.fmrib.ox.ac.uk/fsl). Analyses were performed using a mixed-effects model (FLAME1) and Z statistic images were thresholded using clusters determined by Z>2.58 and a (corrected) cluster significance threshold of P=0.05.*

#### Figure S5: Thumb activity – individual peaks

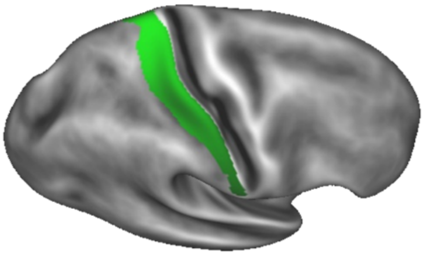

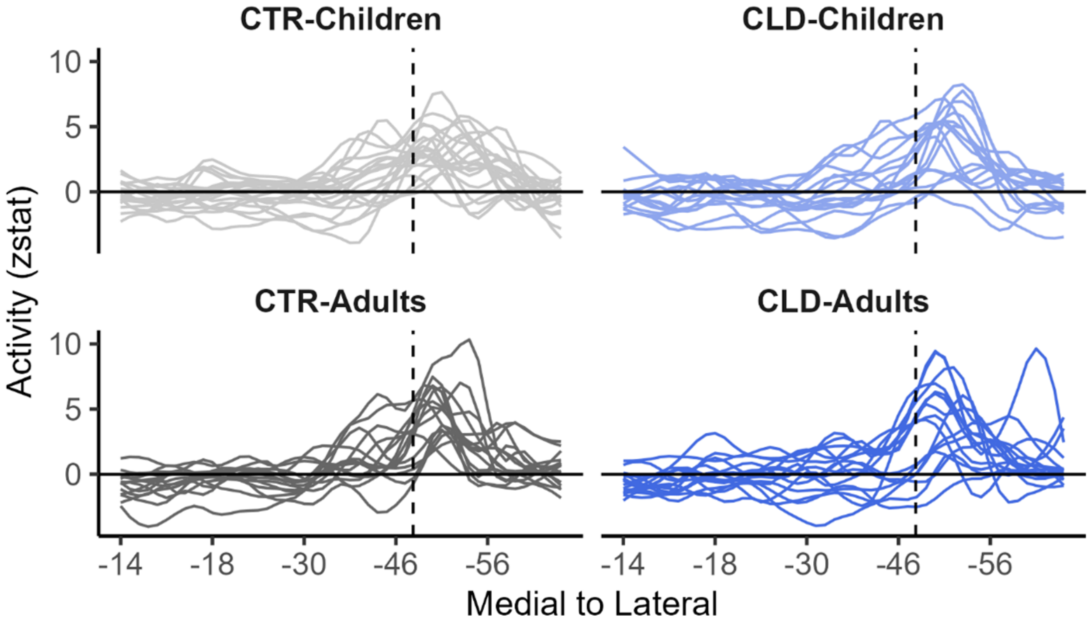


***Figure S5. Individual activity profile for thumb stimulation.*** *Activity (z values) profile plots of all participants along the BA1-2 strip (highlighted in green) within the postcentral gyrus, in the hemisphere ipsilateral to the missing or non-dominant hand (i.e. contralateral to the stimulated thumb). Across all participants, a clear peak of neural activity is observed laterally to the centre of the hand area, which is marked by the vertical dashed line. This region corresponds to the centre of gravity for the five digits, as estimated by a meta-analysis (Holmes & Tamè, 2019). Notably, in one of the CLD adults, the activity profile exhibits a double peak. These findings suggest consistent neural representation of the hand across participants.*

#### Figure S6: Group contrast – CLD > CRT, Arm stimulation

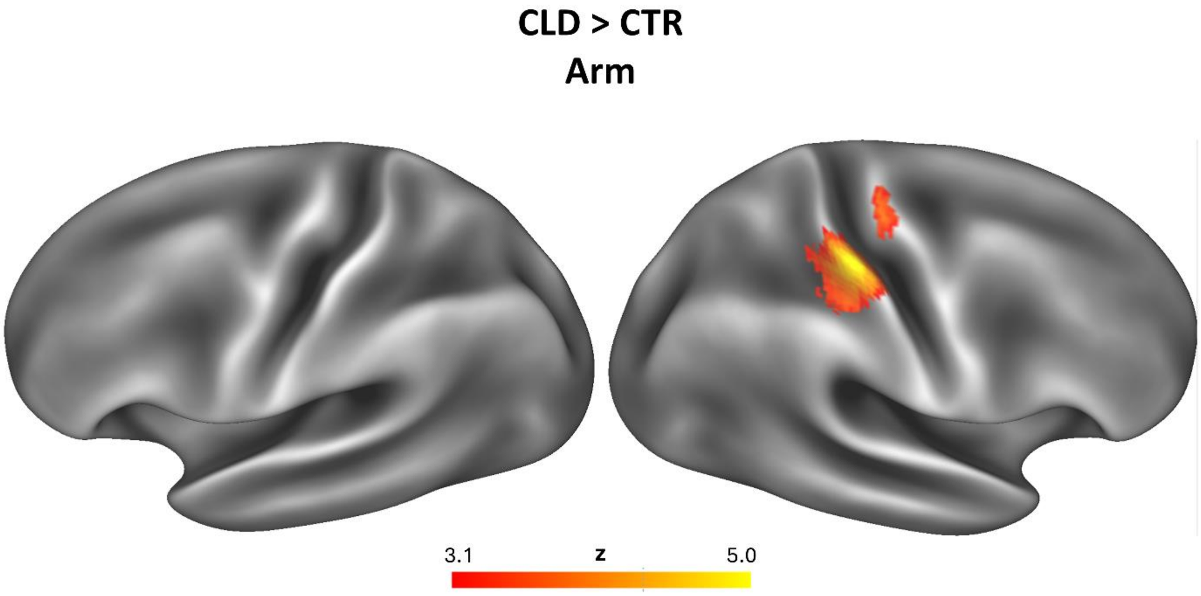


***Figure S6. Differential activity (CLD > CRT) when the Arm is stimulated.*** *Group-contrast (CLD > CTR) activation maps during stimulation of the residual/nondominant arm. Compared to control (CTR) participants, individuals in the CLD group demonstrated significantly greater activation in the contralateral missing-hand representation area. Analyses were performed using a mixed-effects model (FLAME1), with Z-statistic images thresholded at Z > 2.58 and a cluster significance threshold (corrected) of P = 0.05. Colour bar indicates Z values.*

#### Figure S7: Activity profile within BA3b

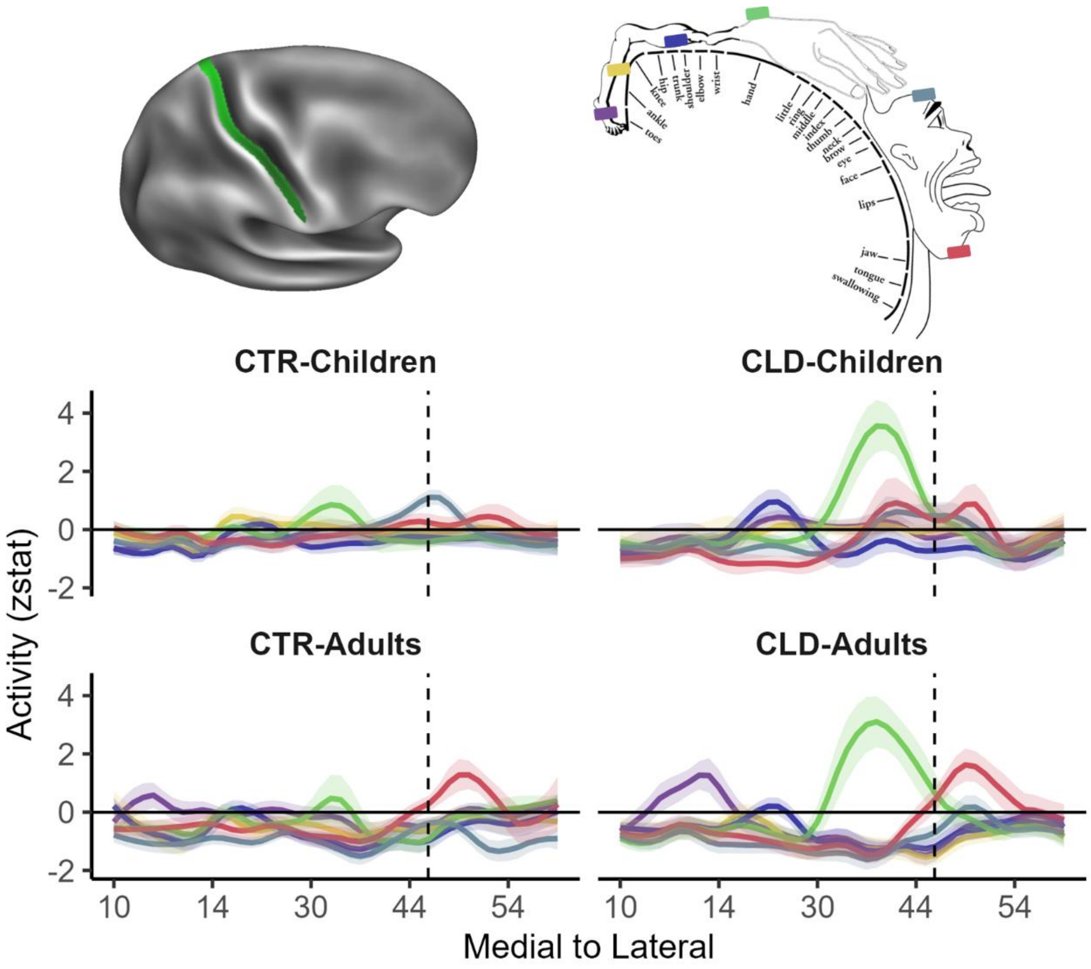


***Figure S7. Global topography: descriptive plots showing remapping across the entire S1 homunculus (area 3b).*** *This plot displays the averaged activity (z-values) profiles for each group and body part along area 3b of S1, providing a comparison with the analyses presented in Figures 4 and 5 of the main manuscript, which focused on the more posterior areas of S1 (i.e., areas 1 and 2). The activity profiles observed in area 3b reveal both similarities and differences compared to those seen in areas BA1-2. Notably, the topographic organization appears more pronounced in this BA1-2 region, particularly for medial activities. For instance, stimulation of lower body parts (foot, leg, and torso) does not consistently activate area 3b, except in the case of the foot for CLD adults, and potentially CRT adults. Additionally, stimulation of the lower face elicits clear activity in area 3b, but predominantly in adults. All annotations and labels in this figure adhere to the same conventions used in Figure 4 for ease of comparison.*

#### Figure S8: Global S1peak activity results.

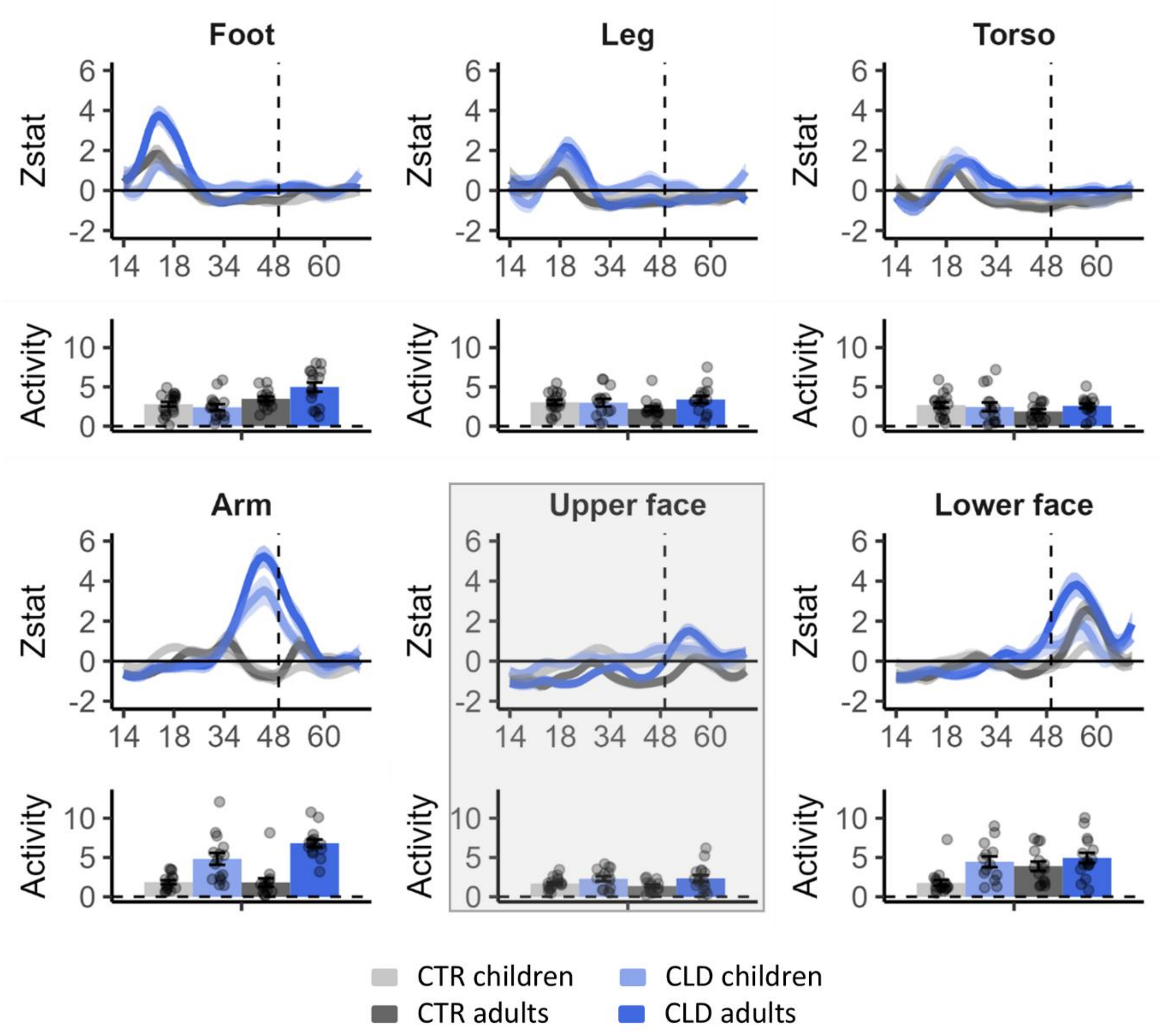


***Figure S8. Averaged activity profiles and peak activities along S1.*** *Averaged activity (z values) profile plots (top panels) along S1 in response to stimulation of different body parts for each group, and the corresponding peak activity values of each individual and related group means (bottom panels). This analysis (detailed in the Supplementary Results section) complements the peak activity shift results presented in Figure 5 of the main manuscript. Notably, stronger activity in CLD participants was observed primarily in body parts adjacent to the deprived hand, specifically the residual arm and lower face, while the upper face remained unaffected. This localised pattern contrasts with the findings from the peak shift analysis (Figure 5), which demonstrated a more widespread impact of sensory deprivation. All annotations and labels in this figure follow the same conventions as in Figure 5.*

#### Figure S9: Initial weights for the main model

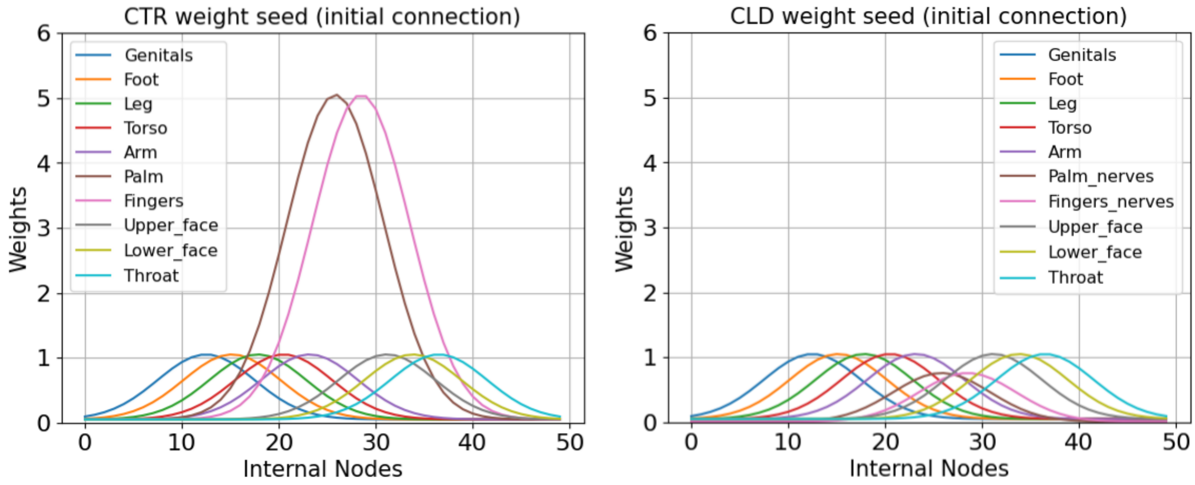


***Figure S9. Initial connection weights between body parts and internal neurons (main model).*** *The figure shows how strongly individual body parts are connected to individual neurons. The left panel represents connections of the CTR group, while the right panel represents connections of the CLD group. For example, neuron 10 has connection with value ~1 with genitalia and connection with value ~0.6 with leg. Notice that for the CLD model (right), palm and fingers are replaced by the corresponding residual nerves and these connections are weaker than the connections of other body parts (see the Results section in the main manuscript). Except for the residual nerves, all bell-shaped curves are vertically shifted by 0.05.*

### Figure S10: Model analysis – Higher frequency usage

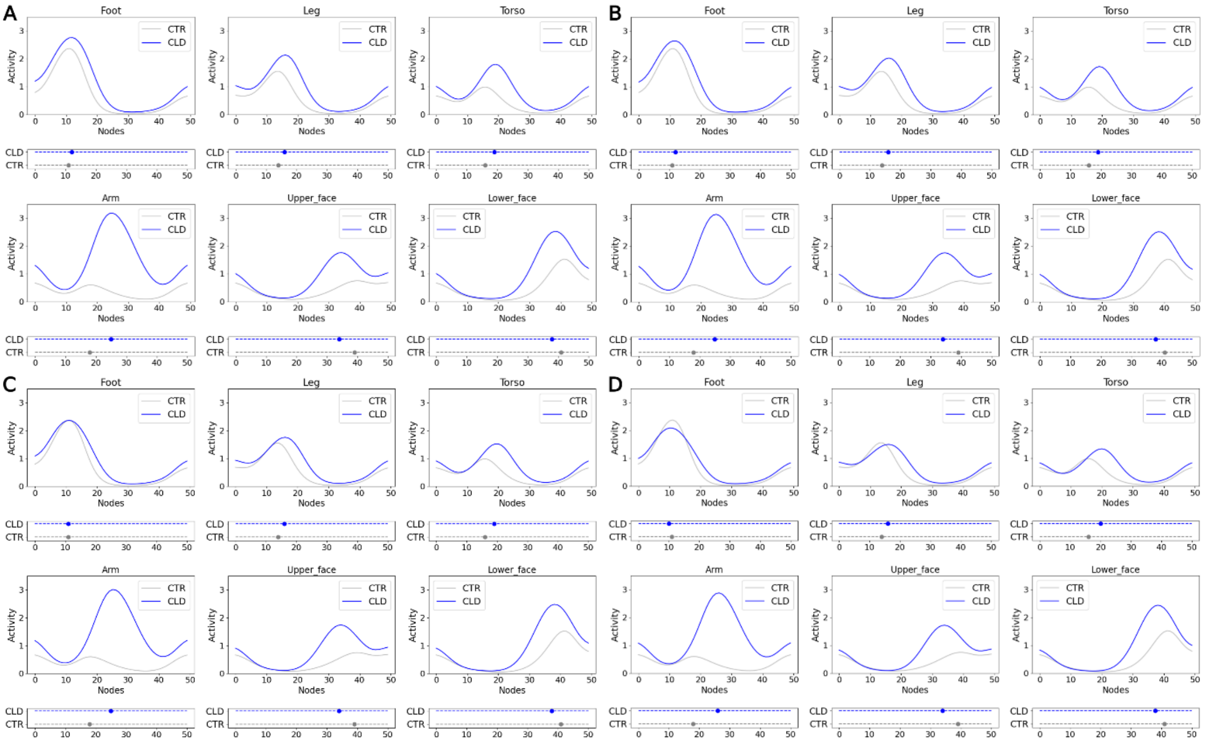


***Figure S10. Effect of higher frequency usage of leg of CLDs on homunculus adaptation.*** *For CTR and CLD activations in subfigure A, baseline parameters are used. For CLD in subfigures B, C and D original stimulation frequency of leg was increased by 25%, 100% and 200%, respectively. Annotations as indicated in Figure 6 of the main manuscript.*

### Figure S11: Model analysis – Lower frequency usage

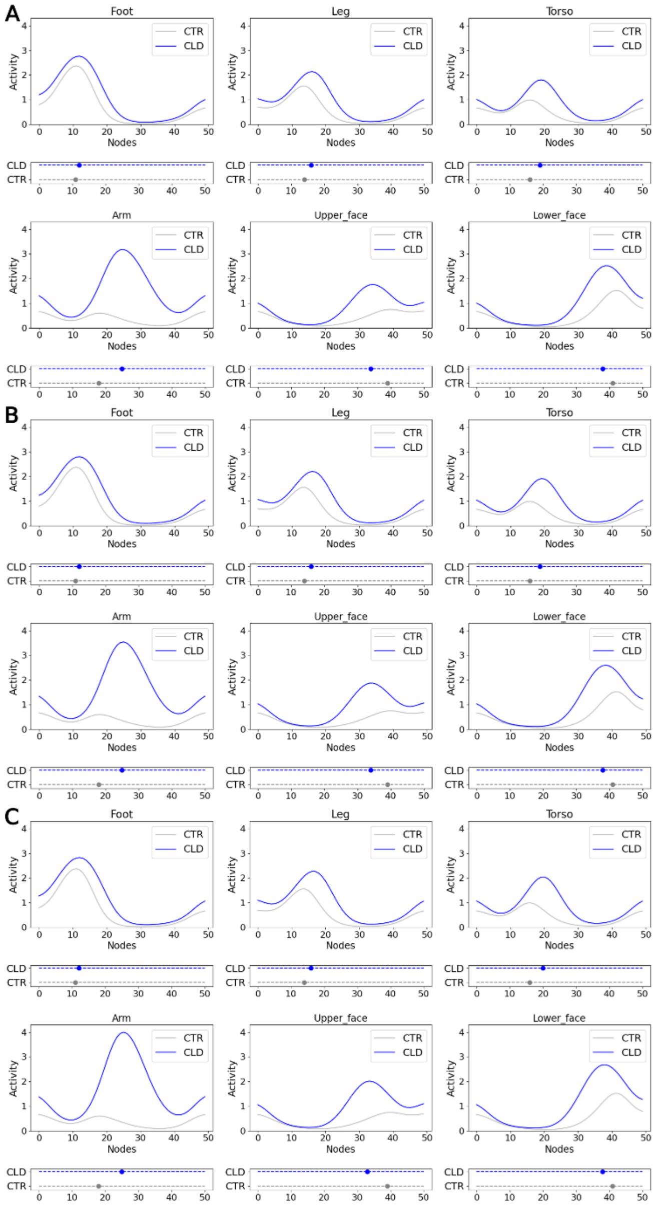


***Figure S11. Effect of lower frequency usage of arm and hand nerves of CLDs on homunculus adaptation.*** *For CTR and CLD activations in subfigure A, baseline parameters are used. For CLD in subfigures B and C, original stimulation frequency of arm and hand residual nerves was decreased by 15% and 30%, respectively. Annotations as indicated in Figure 6 of the main manuscript.*

#### Figure S12: Initial weights for the rudimentary model

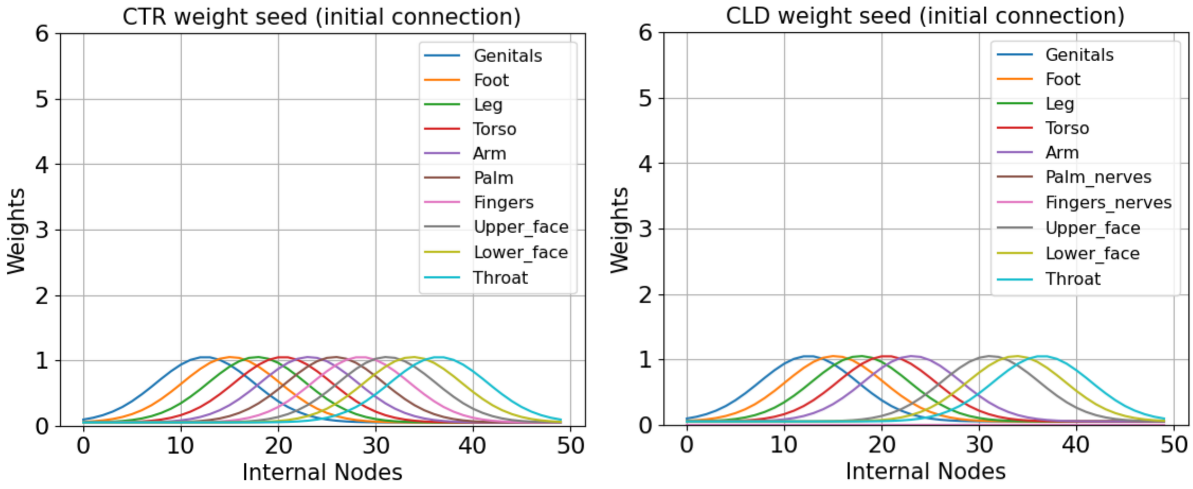


*Figure S12. **Initial connection weights between body parts and internal neurons (rudimentary model)**. (Left) Initial connection weights for CTR group. All body parts have same connection pattern. (Right) Initial connection weights for CLD group. Notice that Hand (i.e., Palm and Fingers) inputs are completely missing. Annotations as indicated in Figure S9.*

#### Figure S13: Rudimentary model results

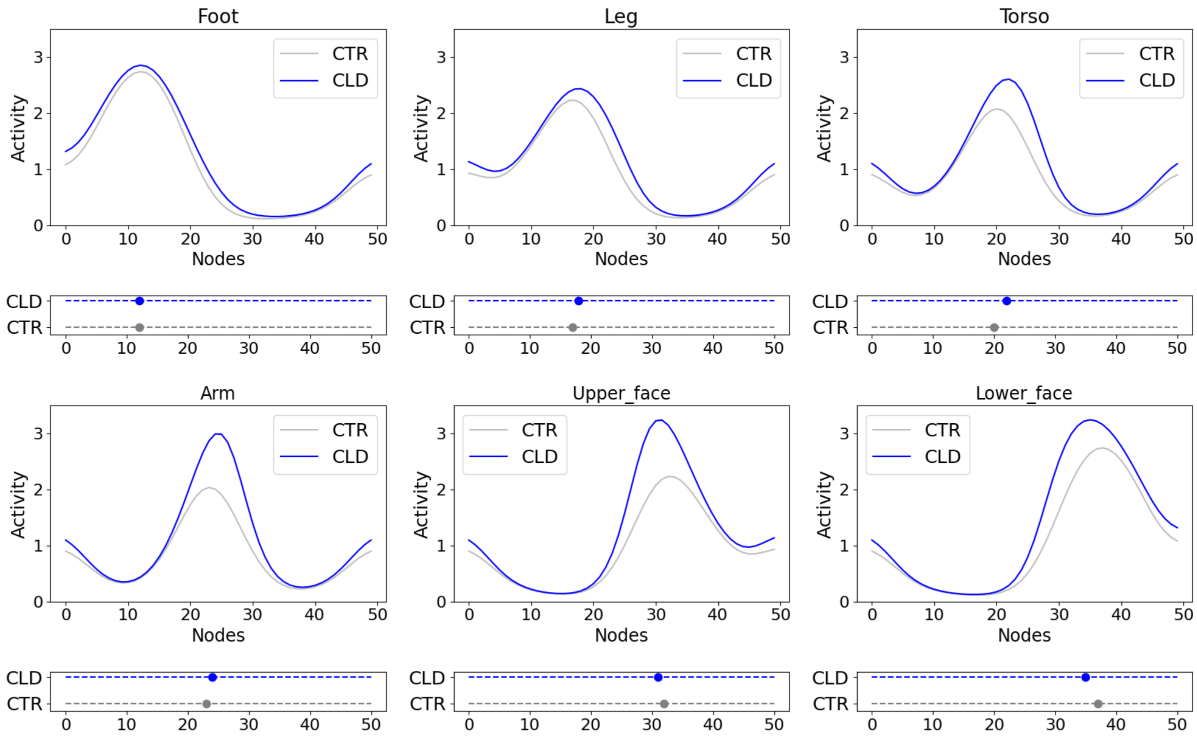


***Figure S13. Peak position and activity plot of the rudimentary model.*** *See text for details. All annotations and labels in this figure follow the same conventions as in Figure 6.*

#### Figure S14: Local remapping beyond S1

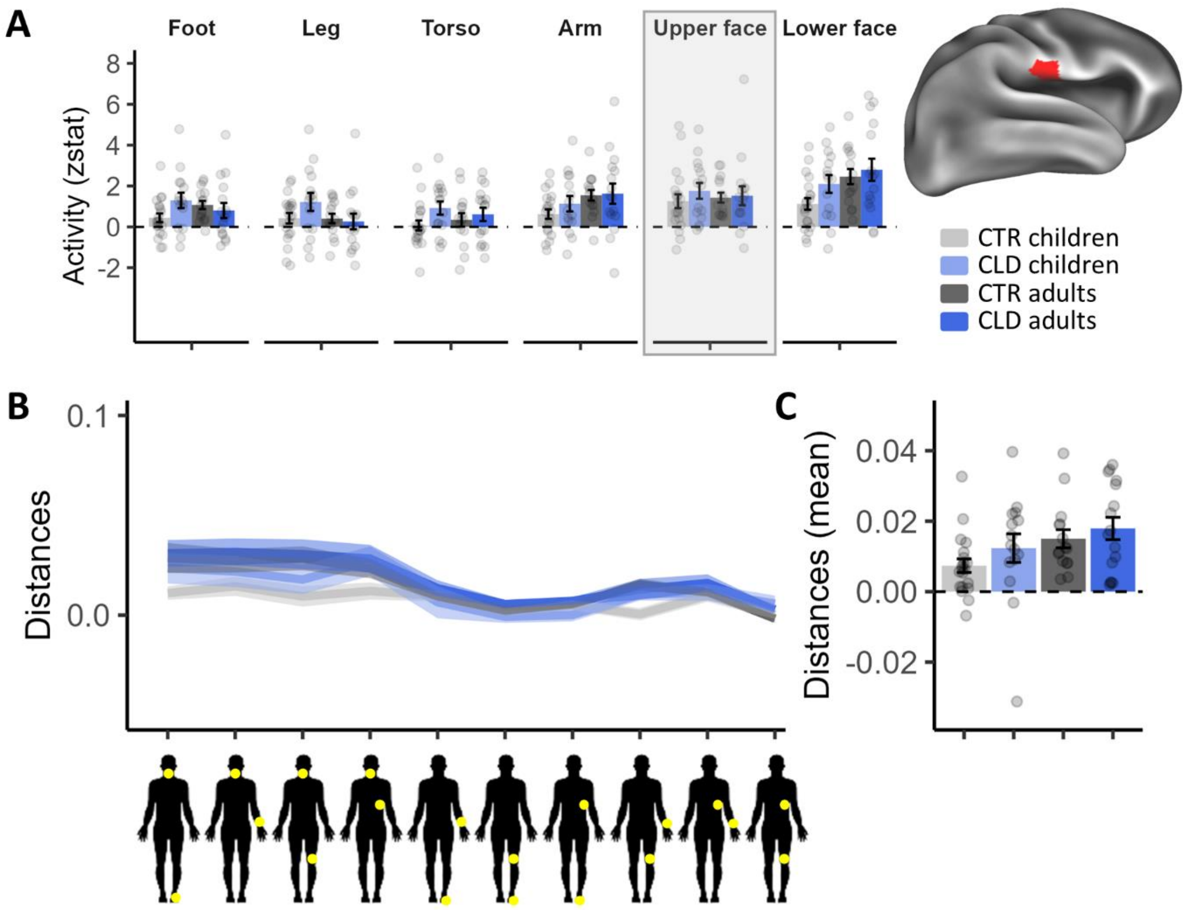


***Figure S14. S2: Activity and Distances show no differential activity due to a congenital limb difference.*** *To investigate topographical changes beyond S1, we defined a ROI within S2 (Panel A, brain plot to the right) using the Op4 label from Glasser's multimodal parcellation (Glasser et al., 2016) in the hemisphere contralateral to the missing/nondominant hand (in CLD/CTR, respectively).* ***(A)*** *Bars indicate z-stat activity within the S2 mask averaged across participants for each group and body part. We found no consistent effects of limb differences which were independent of age.* ***(B-C)*** *Line plot showing pairwise Euclidian distances (RSA analysis) for each group, which were averaged per group. We found a significant main effect for age and no main effect for limb difference nor an interaction. All annotations are as in Figure 3.*

### Supplementary tables

#### Table S1: Behavioural sub-tasks and objects

1. **Padlock:** Put key in the padlock, twist, and remove from suitcase.
2. **Velcro:** Open a Velcro book bag (long Velcro strip).
3. **Plastic envelope:** Open a plastic envelope and remove plaster packet.
4. **Plaster:** Open plaster
5. **Buttons:** Do three buttons of varying sizes (large, medium, small)
6. **Jar:** Remove lid from a jar (unscrew)
7. **Pen:** Remove lid from a pen (pull)
8. **Lego:** Separate Lego bricks, two standard Lego bricks and two flat Lego bricks
9. **Tin:** Remove lid from a wide tin (pull)
10. **Nut and bolt:** Unscrew nut from a small wooden screw (traditional construction toy)
11. **Screwdriver:** Use a screwdriver to put a screw into a wooden block (traditional construction toy)
12. **Glove:** Place a glove on the intact hand
13. **Salad box:** Open a salad box
14. **Beads:** Thread three beads
15. **Sweet:** Open a twisty sweet

#### Linear mixed-models outputs (ANOVA tables)

***Table S2-1: ANOVA output for co-use index***

| Terms | Sum Sq | Mean Sq | NumDF | DenDF | F value | Pr(>F) |
|---|---|---|---|---|---|---|
| Age | 12.10 | 12.10 | 1 | 60.03 | 49.95 | < 0.001*** |
| Limb difference (LD) | 36.90 | 36.90 | 1 | 60.04 | 152.36 | < 0.001*** |
| Age:LD | 1.96 | 1.96 | 1 | 60.04 | 8.08 | 0.006** |

***Table S2-2: ANOVA output for proportion of use time***

| Terms | Sum Sq | Mean Sq | NumDF | DenDF | F value | Pr(>F) |
|---|---|---|---|---|---|---|
| Age | 1.34 | 1.34 | 1 | 60.08 | 34.31 | < 0.001*** |
| Limb difference (LD) | 2.65 | 2.65 | 1 | 60.08 | 67.89 | < 0.001*** |
| Body part (BP) | 72.59 | 18.15 | 4 | 55.99 | 464.26 | < 0.001*** |
| Age:LD | 0.25 | 0.25 | 1 | 60.08 | 6.28 | 0.015* |
| Age:BP | 1.97 | 0.49 | 4 | 4,625.22 | 12.61 | < 0.001*** |
| BP:LD | 24.48 | 6.12 | 4 | 4,625.22 | 156.58 | < 0.001*** |
| Age:LD:BP | 1.00 | 0.25 | 4 | 4,625.22 | 6.41 | < 0.001*** |

***Table S2-3: ANOVA output for thumb peak position analysis***

| Terms | num Df | den Df | MSE | F | ges | Pr(>F) |
|---|---|---|---|---|---|---|
| Age | 1 | 62 | 9.93 | 0.07 | 0.00 | 0.791 |
| Limb difference (LD) | 1 | 62 | 9.93 | 2.96 | 0.05 | 0.091# |
| Age:LD | 1 | 62 | 9.93 | 0.00 | 0.00 | 0.992 |

***Table S2-4: ANOVA output for thumb peak activity analysis***

| Terms | num Df | den Df | MSE | F | ges | Pr(>F) |
|---|---|---|---|---|---|---|
| Age | 1 | 62 | 5.1 | 2.64 | 0.04 | 0.110 |
| Limb difference (LD) | 1 | 62 | 5.1 | 0.03 | 0.00 | 0.859 |
| Age:LD | 1 | 62 | 5.1 | 0.38 | 0.01 | 0.540 |

***Table S2-5: ANOVA output for local remapping activity analysis in the deprived hand area***

| Terms | Sum Sq | Mean Sq | NumDF | DenDF | F value | Pr(>F) |
|---|---|---|---|---|---|---|
| Age | 0.33 | 0.33 | 1 | 62 | 0.30 | 0.586 |
| Limb difference (LD) | 37.32 | 37.32 | 1 | 62 | 34.26 | < 0.001*** |
| Body part (BP) | 6.39 | 1.60 | 4 | 0 | 1.47 | 1.000 |
| Age:LD | 0.94 | 0.94 | 1 | 62 | 0.86 | 0.357 |
| Age:BP | 16.26 | 4.07 | 4 | 248 | 3.73 | 0.006** |
| LD:BP | 108.39 | 27.10 | 4 | 248 | 24.87 | < 0.001*** |
| Age:LD:BP | 8.76 | 2.19 | 4 | 248 | 2.01 | 0.094# |

***Table S2-6: ANOVA output for local remapping distances analysis in deprived hand area***

| Terms | Sum Sq | Mean Sq | NumDF | DenDF | F value | Pr(>F) |
|---|---|---|---|---|---|---|
| Age | 0.02 | 0.02 | 1 | 62 | 10.78 | 0.002** |
| Limb difference (LD) | 0.04 | 0.04 | 1 | 62 | 28.33 | < 0.001*** |
| Age:LD | 0.00 | 0.00 | 1 | 62 | 2.81 | 0.099# |

*Table S2-7: ANOVA output for global peak position analysis in S1*

| Terms | Sum Sq | Mean Sq | NumDF | DenDF | F value | Pr(>F) |
|---|---|---|---|---|---|---|
| Age | 7.26 | 7.26 | 1 | 58.53 | 0.91 | 0.343 |
| Limb difference (LD) | 244.47 | 244.47 | 1 | 58.50 | 30.78 | < 0.001*** |
| Body part (BP) | 21.52 | 5.38 | 4 | 218.43 | 0.68 | 0.608 |
| Age:LD | 31.54 | 31.54 | 1 | 58.49 | 3.97 | 0.051# |
| Age:BP | 40.44 | 10.11 | 4 | 223.10 | 1.27 | 0.281 |
| LD:BP | 867.47 | 216.87 | 4 | 223.06 | 27.31 | < 0.001*** |
| Age:LD:BP | 33.86 | 8.46 | 4 | 223.05 | 1.07 | 0.374 |

*Table S2-8: ANOVA output for global peak activity analysis in S1*

| Terms | Sum Sq | Mean Sq | NumDF | DenDF | F value | Pr(>F) |
|---|---|---|---|---|---|---|
| Age | 10.26 | 10.26 | 1 | 57.88 | 4.29 | 0.043* |
| Limb difference (LD) | 51.15 | 51.15 | 1 | 57.86 | 21.38 | < 0.001*** |
| Body part (BP) | 90.75 | 22.69 | 4 | 221.13 | 9.48 | < 0.001*** |
| Age:LD | 4.11 | 4.11 | 1 | 57.86 | 1.72 | 0.195 |
| Age:BP | 47.49 | 11.87 | 4 | 220.99 | 4.96 | < 0.001*** |
| LD:BP | 143.46 | 35.86 | 4 | 220.96 | 14.99 | < 0.001*** |
| Age:LD:BP | 25.10 | 6.27 | 4 | 220.96 | 2.62 | 0.036* |

***Table S2-9: ANOVA output for local remapping activity analysis in S2***

| Terms | Sum Sq | Mean Sq | NumDF | DenDF | F value | Pr(>F) |
|---|---|---|---|---|---|---|
| Age | 0.33 | 0.33 | 1 | 62 | 0.30 | 0.586 |
| Limb difference (LD) | 37.32 | 37.32 | 1 | 62 | 34.26 | < 0.001*** |
| Body part (BP) | 6.39 | 1.60 | 4 | 0 | 1.47 | 1.000 |
| Age:LD | 0.94 | 0.94 | 1 | 62 | 0.86 | 0.357 |
| Age:BP | 16.26 | 4.07 | 4 | 248 | 3.73 | 0.006** |
| LD:BP | 108.39 | 27.10 | 4 | 248 | 24.87 | < 0.001*** |
| Age:LD:BP | 8.76 | 2.19 | 4 | 248 | 2.01 | 0.094# |

***Table S2-10: ANOVA output for local remapping distances analysis in S2***

| Terms | Sum Sq | Mean Sq | NumDF | DenDF | F value | Pr(>F) |
|---|---|---|---|---|---|---|
| Age | 0 | 0 | 1 | 62 | 5.18 | 0.026* |
| Limb difference (LD) | 0 | 0 | 1 | 62 | 1.87 | 0.177 |
| Age:LD | 0 | 0 | 1 | 62 | 0.12 | 0.725 |

#### Table S3: Double peaks and no peaks

| Group | Reason | Body part | N |
|---|---|---|---|
| CTR Adults | Double peak | Arm | 7 |
| CTR Children | Double peak | Arm | 2 |
| CLD Adults | Double peak | Foot | 1 |
| CTR Adults | Double peak | Foot | 1 |
| CLD Children | Double peak | Leg | 1 |
| CTR Adults | Double peak | Leg | 4 |
| CTR Children | Double peak | Leg | 1 |
| CTR Adults | Double peak | Torso | 3 |
| CTR Children | No peak | Torso | 2 |

#### Table S4: Model specifications

***Table S4:*** *Summary of parameter differences between individual model scenarios. See Methods for details.*

***BP****=body parts,* ***PR****=peripheral reorganization*

| Model | Peripheral Reorganization of CLD | BP stimulation frequency | Initial connection patterns between the BP and nodes |
|---|---|---|---|
| PR + homeostasis | Yes | P=0.15 for palm and fingers and corresponding nerves; P=0.05 for other body parts | CTR: higher magnitude for hand and fingers, CLD: lower magnitude for palm and fingers residual nerves; Both: same magnitude for other BP (see Figure S7) |
| Rudimentary homeostasis | No | P=0.05 for all BP | No connections for palm and fingers for CLD, same magnitude for others (see Figure S12 in Suppl. Mat.) |
| PR + homeostasis + behaviour (CLD leg 25% stimulation increase) | Yes | P=1.25*0.05=0.0625 for CLD leg; For others see “PR + homeostasis” | see “PR + homeostasis” |
| PR + homeostasis + behaviour (CLD leg 200% stimulation increase) | Yes | P=3*0.05=0.15 for CLD leg; For others see “PR + homeostasis” | see “PR + homeostasis” |
| PR + homeostasis + behaviour (arm) | Yes | 0.7*0.15=0.105<P<0.15 for CLD palm and fingers nerves, 0.7*0.05=0.035<P<0.05 for CLD arm; For others see “PR + homeostasis” | see “PR + homeostasis” |

#### Table S5: Model observations

***Table S5. Alignment of the models with the key empirical observations Observation1-4*** *(see Section "Key observations to be explained by the model" in the Methods section of the main text). Observation 1: Arm peaks difference CLD vs CTR; Observation 2: Activation peak shifts CLD vs CTR; Observation 3: Higher activation peaks magnitudes for CLD; Observation 4: Decreasing activation magnitudes.*

***BP****=body parts,* ***PR****=peripheral reorganization (see Table S4 for summary of individual specifications)*

| Model | Obs. 1 | Obs. 2 | Obs. 3 | Obs. 4 |
|---|---|---|---|---|
| PR + homeostasis | Yes | Yes | Yes | Yes |
| Rudimentary homeostasis | Yes | Yes for all BP except for foot | Yes | No |
| PR + homeostasis + behaviour (CLD leg 25% stimulation increase) | Yes | Yes | Yes | Yes |
| PR + homeostasis + behaviour (CLD leg 200 % stimulation increase) | Yes | No | No | Yes |
| PR + homeostasis + behaviour (arm) | Yes | Yes | Yes | Yes |

#### Table S6: Group contrasts

***Table S5.*** *MNI coordinates, p-value, z score and number of voxels for the group analysis shown in Figure S6 (CLD > CTR, arm stimulation). Analyses were performed using a mixed-effects model (FLAME1), with Z-statistic images thresholded at Z > 2.58 and a cluster significance threshold (corrected) of P = 0.05.*

| | **Contrast** | **Voxels** | **P** | **MAX** | | | | **COG** | | |
|---|---|---|---|---|---|---|---|---|---|---|
| | | | | z | x | y | z | x | y | z |
| CLD > CLD | | | | | | | | | | |
| | Arm | 494 | <0.001 | 4.31 | 48 | -24 | 56 | 47.6 | -23.9 | 55.8 |
